# EXPRESSION INSIGHTS INTO GASTRIC ADENOCARCINOMA: NETWORK ANALYSIS REVEALS KEY HUB GENES AND FUNCTIONAL MODULES

**DOI:** 10.1101/2025.08.27.672661

**Authors:** Ronald Matheus da Silva Mourão, Fabiano Cordeiro Moreira, Jéssica Manoelli Costa da Silva, Daniel de Souza Avelar da Costa, Valéria Cristiane Santos da Silva, Amanda Ferreira Vidal, Leandro Lopes de Magalhães, Ana Karyssa Mendes Anaissi, Samia Demachki, Williams Fernandes Barra, Paulo Pimentel de Assumpção

## Abstract

Gastric cancer remains a leading cause of cancer‐related mortality worldwide, with poor survival rates. To uncover its molecular basis, we performed weighted gene coexpression network analysis on RNA‐seq data from 119 gastric adenocarcinoma (GAC) and peritumoral tissue (PTT) samples. We identified six key coexpression modules: MEmagenta, MEbrown, MEgreen, and MEturquoise showed strong positive correlations with GAC, whereas MEblack and MEblue were negatively correlated. Hub genes such as *COL3A1, PGAM1*, and *CFL1* were among the most highly expressed in GAC samples compared to PTT. ROC analyses of selected hub genes yielded AUC values exceeding 0.80 for distinguishing GAC from PTT, underscoring their diagnostic potential. Integrating cellular deconvolution with module expression revealed that MEblack hub genes (*TFF1, TFF2, GKN1, GKN2* and *MUCL3*) correlated positively with B-cell abundance and negatively with resting mast cells and neutrophils. Conversely, MEturquoise hub genes (*COL1A2, COL3A1, TAGLN* and *SPARC*) correlated strongly with Cance Associated Fibroblasts and inversely with B-cells, reflecting a collagen-rich stroma. We also observed overlapping expression profiles between GAC and PTT, indicating tumor heterogeneity and a molecular continuum between normal and neoplastic tissues. These integrated insights highlight candidate biomarkers and in GAC.

## Introduction

Gastric cancer (GC) is a pressing global health issue, ranking as the fifth most prevalent cancer with significant rates of incidence and mortality affecting both men and women (Bray et al. 2024) Despite a century of progress resulting in declining rates—thanks to early diagnosis and the successful eradication of Helicobacter pylori—GC remains a formidable challenge (Lin et al. 2024). Alarmingly, the current 5-year survival rate for patients is still below 30% (D’Alpino Peixoto et al. 2020)

Understanding the intricacies of gastric cancer is crucial for overcoming therapeutic hurdles. While research has made substantial progress in uncovering its causes, the heterogeneous nature of GC and the complexity of its gene expression profiles hinder effective treatment(Joshi and Badgwell 2021) A deep and comprehensive understanding of the underlying molecular mechanisms is essential to combat this.

Recent advancements in systems biology are transforming our approach to GC. The construction of weighted gene coexpression networks offers an invaluable tool for identifying gene modules associated with disease phenotypes. These networks facilitate exploring functional relationships among genes and reveal their collective influence on tumor biology. By analyzing these intricate coexpression patterns, researchers can identify key gene networks that correlate with specific cancer characteristics, thereby uncovering promising therapeutic targets (Li et al. 2021; Zheng et al. 2021; Ding et al. 2023).

This study focuses on constructing and analyzing coexpression networks from gastric adenocarcinoma (GAC) and adjacent peritumoral tissues (PTT). Utilizing a scale-free topology to build these networks has enabled us to identify distinct coexpression patterns and explore their connections to tumor characteristics. This comprehensive approach clarifies how genes interact and emphasizes the critical roles of hub genes within these networks.

The present findings offer significant insights into the gene networks associated with GC, providing potential new avenues for targeted therapies. By elucidating the roles of key hub genes and their associated networks, this study provides a comprehensive framework for understanding the molecular basis of GC. Furthermore, it underscores the potential for novel biomarker discovery and therapeutic strategies.

## Methods

### Sample Characterization and Ethical Aspects

A total of 119 samples of tumor and PTT were collected from patients diagnosed with the most common type of GC, GAC. The subjects were recruited from the João de Barros Barreto University Hospital in Belém, Brazil. The recruitment of participants and the collection of samples for this study occurred from July 2, 2022, to July 6, 2023. Prior to participating in the study, the researchers clearly communicated the objectives to the participants, ensuring they fully understood the purpose of the research. Subsequently, the participants willingly provided their written informed consent. The study was conducted following the Declaration of Helsinki and received approval from the Ethics Committee of the João de Barros Barreto University Hospital (Approval number: 47580121.9.0000.5634).

### Clinical Characterization of Patients

81 patients were the subject of comprehensive medical records review and analysis of ancillary clinical data. The review included the following variables: the presence of GAC, ypTNM, Laurèn classification, MMR status, E-cadherin mutation, and expression of proteins PDL1, PSM2, and MLH1.

### Extraction and Quality of the Total RNA

Approximately, 50-100 mg of tissue from each sample was macerated. Subsequently, 1 ml of TRIZOL® reagent was added to the processed tissue for extraction. Total RNA integrity and concentration were analyzed in the Qubit 4.0 Fluorometer (Thermo Fisher Scientific) and NanoDrop ND-1000 (Thermo Fisher Scientific). The optimal criteria met for total RNA integrity corresponded to values between 1.8 and 2.2 (A260/A280 ratio), >1.8 (A260/A230 ratio), and RIN (RNA integrity number) ≥ 5. The cutoff accounted for RNA quality variability in clinical samples, balancing cohort representation with data reliability.

### Construction of cDNA Libraries and Sequencing

The TruSeq Stranded Total RNA Library Prep Kit with Ribo-Zero Gold (Illumina) was employed to remove cytoplasmic and mitochondrial rRNA. Libraries were then processed using the *NextSeq*® 500 High Output V2 kit - 150 cycles (*Illumina*) under the conditions specified by the manufacturer. After constructing the libraries, a new integrity assessment was performed on the 2200 TapeStation System (Agilent). The previously created cDNA libraries were loaded into the Illumina NextSeq sequencing system (Illumina) and sequenced via pair-end.

### Quality, Alignment, Quantification and Transcriptome Expression

The initial evaluation of sequencing read quality was conducted using FastQC (v0.11.9). To enhance data integrity, low-quality reads and adapter sequences were eliminated using Trimommatic, applying a minimum Phred quality score threshold of QV15. This threshold was chosen considering the robustness of Salmon’s k-mer-based pseudoalignment to moderate variations in base quality. Post-filtering, transcript-level quantification was performed with Salmon (v1.5.2), aligning the reads against the human transcriptome reference (hg38). The resulting transcript abundances were imported via Tximport, facilitating the creation of a DESeq2 object. This object enabled gene expression levels, accounting for variables such as tissue type (GAC or PTT) and sequencing batch effects. For downstream analyses, variance-stabilized and batch-corrected expression values were utilized.

### Co-expression Network Construction and Hub Genes Selection

To construct a gene co-expression network, we used the Weighted Gene Coexpression Network Analysis (WGCNA) package, version 1.73. (Langfelder and Horvath 2008) Genes with expression levels below one read in more than 50% of the samples were excluded from the analysis to reduce the number of genes with low information content, reduce noise, and avoid artificial correlations. We selected 5,000 genes with the highest variance to prioritize those that are biologically relevant and informative ( Table S1 ). This ensures that the analysis captures meaningful co-expression patterns associated with key biological processes. The weighted median-based correlation (Biweight midcorrelation) similarity matrix was computed for gene pairs. A soft threshold power value of β = 12 was selected to create a scale-free network. The first principal component of each module’s gene expression was calculated as Module Eigengenes (MEs), and Spearman correlations between MEs and clinical aspects were assessed. Gene significance (GS) for traits and Module Membership (MM) were determined for each gene ( Table S2 ). The Grey module consisted of genes that did not demonstrate a correlated expression pattern with other genes. The filtration of genes within each module was executed based on the following criteria: |GS| > 0.3, |MM| > 0.6, and p<0.05. We used Cytoscape version 3.7.9 software for visualization, biological interpretation, and identifying hubs.

Hub genes are defined as those with the highest connectivity degrees, playing central roles in the biological network. Gene’s connectivity can provide significant insights into its functional relevance and impact on disease progression. To identify hub genes in this study, the top 100 edges with the highest connectivity degrees within the co-expression networks were selected, and hubs were defined as genes with the highest number of associations with other genes.

### Gene Ontology

To describe biological pathways and functions related to the modules, the genes were subjected to functional enrichment analysis. For this attempt, the Gene Ontology (GO) tool (geneontology.org/) was used via ClusterProfile v4.3.2 (Yu et al. 2012). The enrichment analysis was performed for genes in the modules with a correlation of the absolute values of GS and MM greater than 0.25 ( Table S3 ). We assume that functional enrichment terms with adjusted p-values less than or equal to 0.05 are meaningful.

### ROC Curve

The Receiver Operating Characteristic Curve (ROC) for the expression of each gene was plotted via package pROC v1.18 (Robin et al. 2011). The ROC curve represents the trade-off between sensitivity and specificity across different thresholds, providing insight into the gene’s ability to distinguish between clinical conditions. The Area Under the Curve (AUC) was then calculated to quantify the gene’s diagnostic accuracy, with values closer to 1 indicating better performance ( Table S4 ). An AUC greater than 0.7 is considered acceptable, reflecting the gene’s potential as a clinical biomarker.

### Immune Cell Deconvolution Analysis

To estimate the relative abundance of various immune cell types within bulk RNA-Seq samples, we applied three computational deconvolution methods. Gene expression levels were quantified as Transcripts Per Million (TPM). Cell fractions were estimated using CIBERSORT with the LM22 immune signature matrix, alongside the quanTIseq and EPIC algorithms. Differences in estimated cell proportions between experimental conditions (GAC and PTT) were evaluated using the Wilcoxon rank-sum test, applying Benjamini-Hochberg correction for multiple testing (FDR ≤ 0.05). Furthermore, Spearman correlation analysis explored relationships between significantly different immune cell fractions and hub gene expression levels in samples. Key results were visualized using composition bar plots, box plots, and correlation matrices.

## Results

Initially, we compared the gene expression between tissues: GAC (n = 65) and PTT to GAC (n = 54). The normalized expression of 5,000 most variable coding genes was evaluated in WGCNA. A hierarchical clustering dendrogram of genes was constructed (Figure 1A) after selecting a soft-thresholding power of β = 12 (Figure 1C). Module eigengenes for each identified coexpression module were then correlated with the two tissue types (Figure 1B).

**Figure 1.**
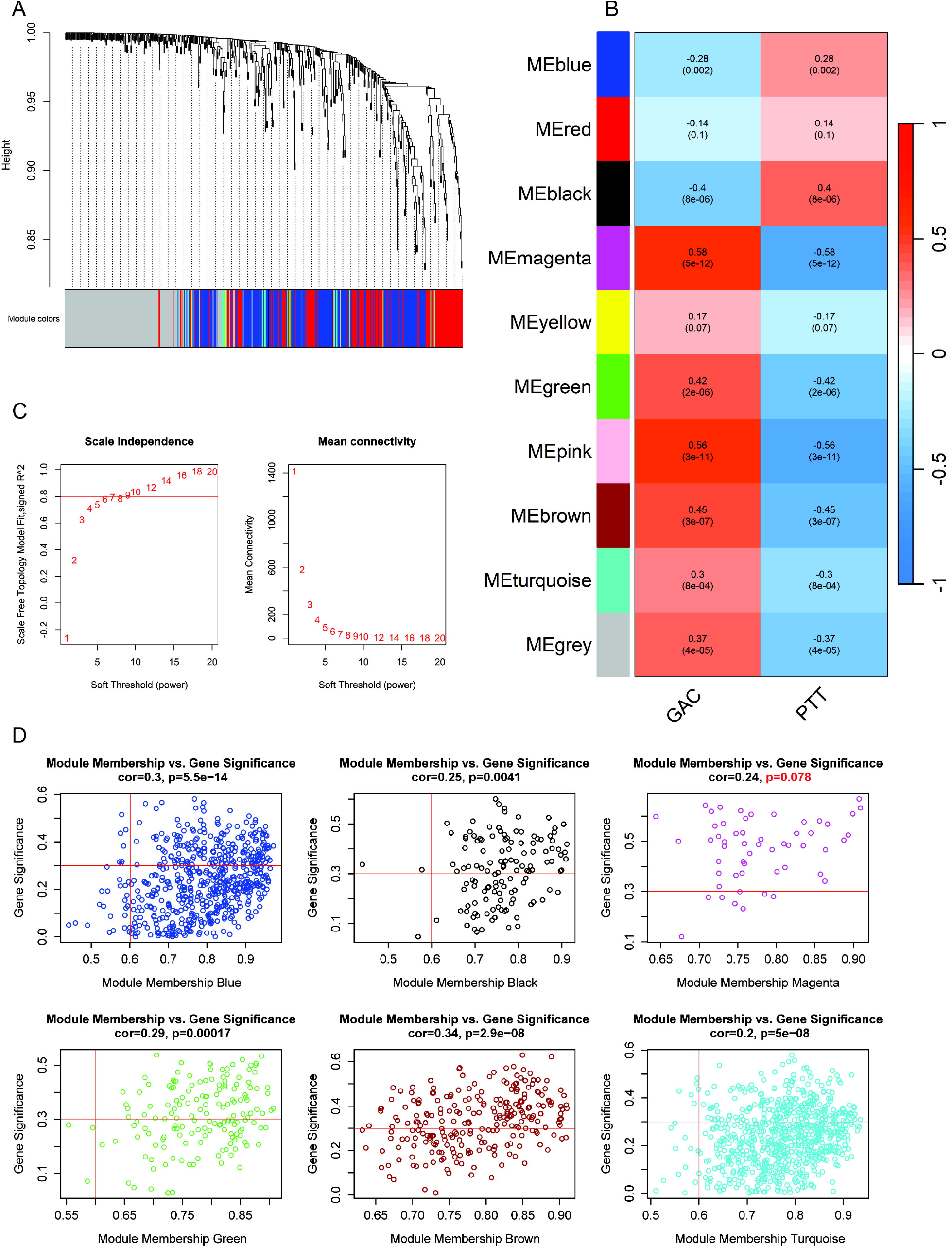
Network construction. (A) Cluster Dendrogram: Hierarchical clustering of genes based on topological overlap to identify distinct coexpression modules. (B) Module–Clinical Relationship: Heatmap showing correlations between module eigengenes and clinical traits. Each cell’s color reflects the strength and direction of the correlation coefficient. (C) Scale Independence: Plot of the scale-free topology fit index (y-axis) versus soft-thresholding power (x-axis), used to select the optimal β for network construction. (D) Gene Significance (GS) vs. Module Membership (MM): Scatter plots depicting the correlation between GS and MM for each key module.

### Key Networks Correlated with GAC or PTT

Following the correlation analysis between the module eigengene values of each co-expression group and tissue status (GAC or PTT), we identified co-expression groups associated with these two tissue types. The MEmagenta (54 genes, r = 0.58, p-value = 5e-12), MEbrown (226 genes, r = 0.45, p-value = 3e−07), MEgreen (145 genes, r = 0.42, p-value = 2e−06), and MEturquoise (487 genes, r = 0.30, p-value = 8e−04) modules were positively correlated (groups called as positive modules) with tumoral tissues (Fig 1c). Although MEpink and MEgrey had r ≥ 0.30 and p-value ≤ 0.05, they did not show significant associations between Gene Significance and Module Membership (Fig 1C), suggesting a lower likelihood that the hub genes in these modules have strong correlations with GAC or PTT. Conversely, the MEblack (105 genes, r = 0.40, p-value = 8e−06) and MEblue (397 genes, r = 0.28, p-value = 0.002) modules were negatively correlated (groups called negative modules) with the expression of GAC (Fig 1B).

### Module Associated with Extracellular Matrix Dynamics

Functional enrichment analysis of gene ontologies was performed to associate the gene groups in each significant module with their potential biological pathways. The MEmagenta (Figure 2A), MEbrown (Figure 2C), MEgreen (Figure 2E), and MEturquoise (Figure 2G) modules were positively associated with tumor-related biological processes in GAC (Figure 2).

**Figure 2.**
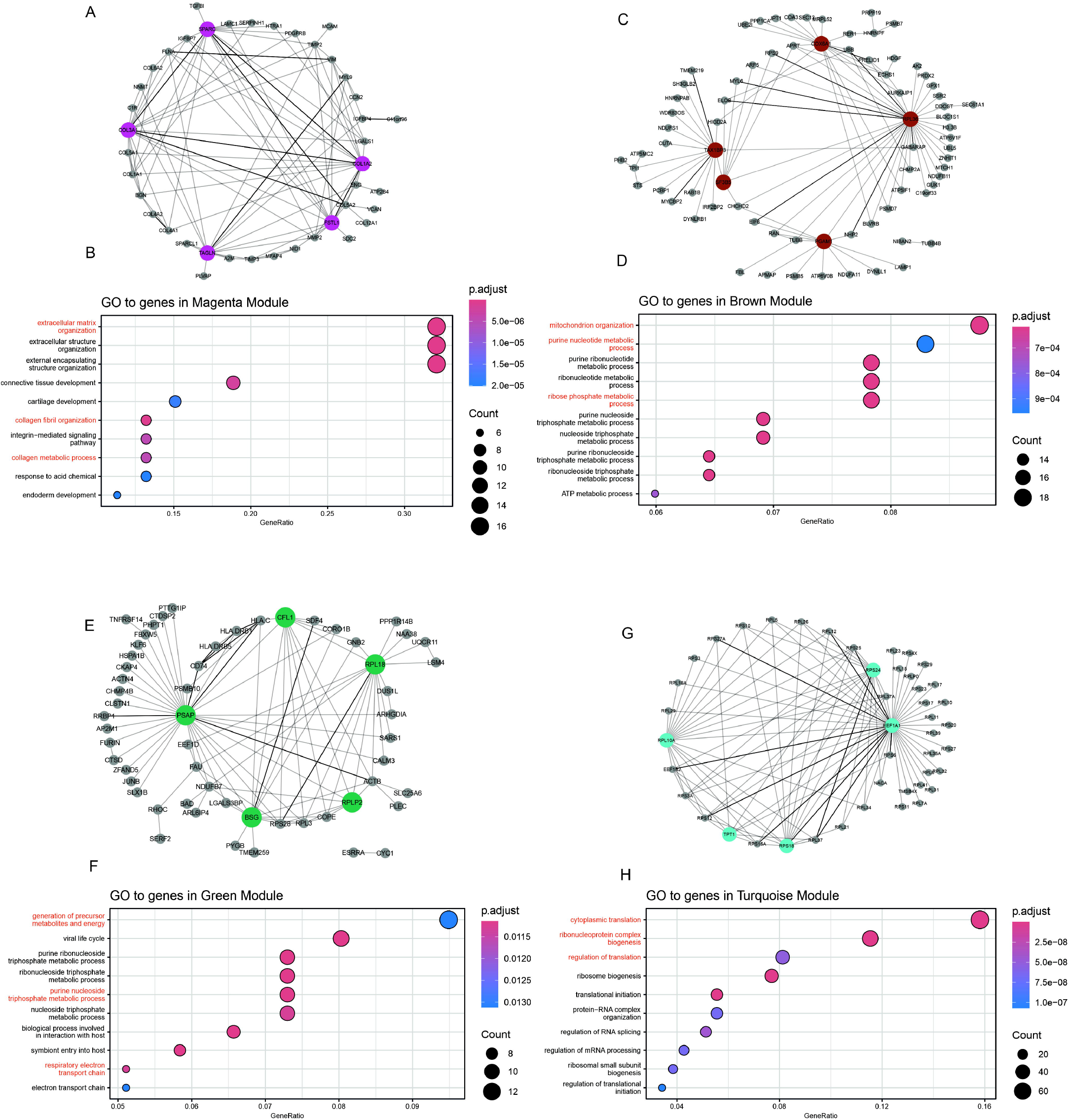
Positively Associated Coexpression Network and Functional Enrichment Analysis. The network of co-expressed genes within the Magenta (A), Brown (C), Green (E), and Turquoise (G) networks is represented. These networks were visualized using the software program Cytoscape. The nodes in the network represent individual genes, while the edges represent the co-expression relationships between them. The accompanying dot plot displays the top ten enriched GO terms associated with module Magenta (B), module Brown (D), module Green (F), and module Turquoise (H).

The most significantly enriched GO terms in the MEmagenta module (Figure 2B), relevant to cancer, included: GO0030198 extracellular matrix organization (gene ratio = 17/53), GO0030199 collagen fibril organization (gene ratio = 7/53), and GO0032963 collagen metabolic process (gene ratio = 7/53). Within the context of networks positively associated with GAC, MEmagenta (min. weight = 0.02 and degree = 41) stood out, with the hub genes being *COL3A1* (17 degrees), *SPARC* (16 degrees), *COL1A2* (16 degrees), *FSTL1* (15 degrees), and *TAGLN* (14 degrees) (Figure 2A).

### Module Associated with Mitochondrial Organization and Purine Metabolism

In the MEbrown module, the most relevant GO terms (Figure 2D) were GO:0007005 mitochondrion organization (gene ratio = 19/217), GO0006163 purine nucleotide metabolic process (gene ratio = 18/217), and GO0019693 ribose phosphate metabolic process (gene ratio = 17/217). Among the networks positively associated with GAC, MEbrown (min. weight = 0.05 and nodes = 75) featured hub genes *RPL36* (34 degrees), *COX6A1* (17 degrees), TAX1BP3 (16 degrees), *PGAM1* (12 degrees), and *SF3B5* (8 degrees) (Figure 2C)

### Module Associated with Energy Production and Purine Metabolism

To the MEgreen module, the most prominent GO terms (Figure 2F) were GO0006091 generation of precursor metabolites and energy (gene ratio = 13/137), GO0009144 purine nucleoside triphosphate metabolic process (gene ratio = 10/137), and GO0022904 respiratory electron transport chain (gene ratio = 7/137). This module, MEgreen (min.weight = 0.05 and degree = 48), featured hub genes *PSAP* (34 degrees), *RPL18* (17 degrees), *BSG* (14 degrees), *CFL1* (12 degrees), and *RPLP2* (8 degrees) (Figure 2E).

### Module Associated with Cytoplasmic Translation and Ribonucleoprotein Biogenesis

The MEturquoise module reveals significant Gene Ontology (GO) terms (Figure 2H): GO0002181 cytoplasmic translation (gene ratio = 74/468), GO0022613 ribonucleoprotein complex biogenesis (gene ratio = 54/468), and GO0006417 regulation of translation (gene ratio = 38/468). This module, MEturquoise (minimum weight = 0.14 and number of nodes = 45), includes hub genes such as EEF1A1 (44 degrees), RPL10A (19 degrees), RPS24 (17 degrees), RPS18 (14 degrees), and TPT1 (8 degrees) (Figure 2G).

### Module Associated with Digestion and Gastrointestinal Epithelium Maintenance

MEblack module GO terms (Figure 3C) included GO0007586 digestion (gene ratio = 14/98), GO0009410 response to xenobiotic stimulus (gene ratio = 15/98), and GO0030277 maintenance of gastrointestinal epithelium (gene ratio = 04/98). MEblack (min.weight = 0.12 and nodes = 29) featured hub genes *GKN2* (16 degrees), *TFF2* (14 degrees), *MUCL3* (14 degrees), *TFF1* (13 degrees), and *GKN1* (13 degrees) (Figure 3C).

**Figure 3.**
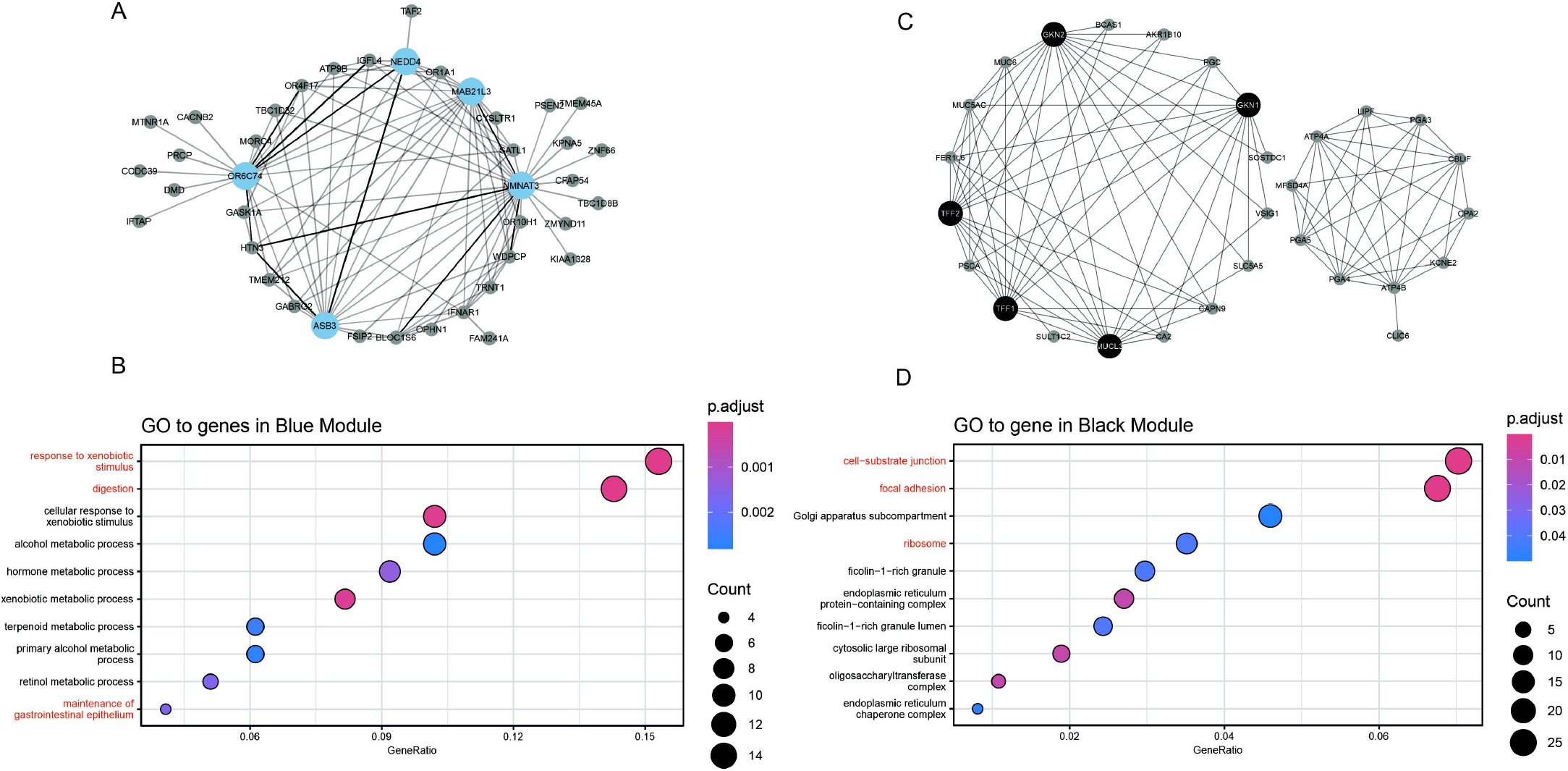
Negative Coexpression Network and Functional Enrichment Analysis. Visualization of the (A) Blue and (C) Black co-expression networks, highlighting the most interconnected hub genes. The adjacent plot shows the enriched pathways identified through Cytoscape, indicating the potential biological functions associated with the module. The dot plot illustrates the top 10 enriched GO terms associated with the (B) Blue and (D) Black networks.

### Module Associated with Cell Adhesion and Ribosome Function

The MEblue module displayed relevant GO terms (Figure 3B) such as GO0030055 cell-substrate junction (gene ratio = 26/370), GO0005925 focal adhesion (gene ratio = 25/370), and GO0005840 ribosome (gene ratio = 13/370). MEblue (min.weight = 0.26 and nodes = 40) had hub genes *NMNAT3* (27 degrees), *OR6C74* (19 degrees), *MAB21L3* (18 degrees), *ASB3* (17 degrees), and *NEDD4* (10 degrees) (Figure 3A).

### Transcript Expression of Gene Hubs and Correlation with Clinical Features

After normalization, characteristic expression patterns were observed for each module. MEbrown, MEgreen, MEmagenta, and METurquoise appear to have similar expression patterns. The hubs in MEblack and MEblue display distinct patterns (Figure 4A).

**Figure 4.**
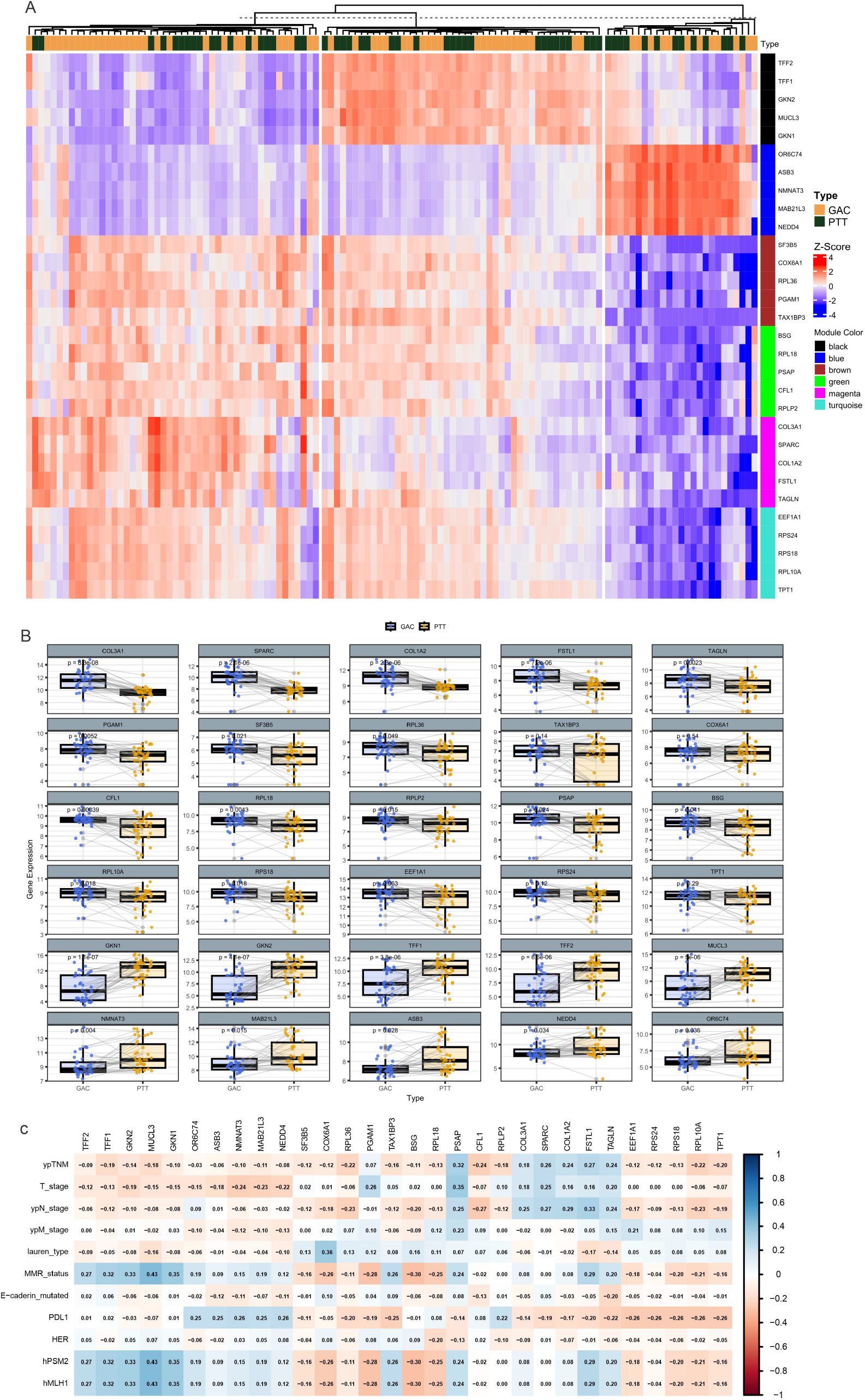
Heatmap of hub genes and paired mean comparison. (A) Heatmap displaying gene expressions of hub genes across all study samples, with rows representing genes and columns representing individual samples. (B) Paired comparison of mean gene expression of these hub genes between GAC tissues and PGAC tissues, highlighting significant differences in expression profiles.(C) Correlation between hub genes and clinical features of patients.

Hierarchical clustering and PCA (Fig. 4A, S1 Fig. ) revealed overlapping expression patterns between GAC and PTT. To explore this further, we conducted a paired expression analysis across modules (Fig. 4B) and compared the expression changes with the AUC values. In the MEmagenta module, *COL3A1, SPARC, COL1A2, FSTL1*, and *TAGLN* were significantly overexpressed in GAC, with a mean AUC of 0.82, supporting their relevance in tumor-associated matrix remodeling. A similar pattern was observed in the MEbrown module for *PGAM1, SF3B5*, and *RPL36* (mean AUC = 0.73), although *COX6A1* showed only comparable expression with a lower AUC of 0.64. All hub genes in the MEgreen module - *CFL1, RPL18, PSAP, BSG*, and *RPLP2* - were also overexpressed in GAC (mean AUC = 0.74). In MEturquoise, *RPL10A, RPS18, EEF1A1*, and *RPS24* were moderately upregulated (mean AUC = 0.68), while *TPT1* showed a weaker association (AUC = 0.62). In contrast, the MEblue genes *NMNAT3, ASB3, OR6C74, NEDD4*, and *MAB21L3* were consistently downregulated in GAC (mean AUC = 0.75). Similarly, MEblack genes *GKN1, GKN2, MUCL3, TFF1*, and *TFF2* showed marked reduction in expression (mean AUC = 0.76), indicating a loss of this mucosal signature in GAC.

The analysis of hub gene expression revealed distinct patterns of correlation with clinically relevant variables in GAC (Fig. 4C). Negative correlations were observed between *PGAM1* (r = −0.25), *BSG* (r = −0.24), *RPL18* (r = −0.23), and *RPS24* (r = −0.21) and the ypTNM stage. In addition, *RPL18* also showed a negative correlation with T stage (r = −0.22). In contrast, positive correlations were identified for *PGAM1* (r = 0.22) and *BSG* (r = 0.21) with the ypN. Regarding Lauren classification, *TFF1* showed a positive correlation (r = 0.21), while *TFF2* and *GKN1* each correlated at r = 0.20. Notably, *TFF1, TFF2, GKN1*, and *GKN2* showed positive correlations with MMR status, hPSM2, and hMLH1, with r values ranging from 0.27 to 0.43. Furthermore, expressions of PDL1 were positively correlated with *OR6C74, ASB3, NMNAT3*, and *MAB21L3* (r = 0.25), while negative correlations were observed between PDL1 and *RPS24, RPS18*, and *RPL10A* (r = −0.26).

### Cellular Composition and Correlation with Hub Genes

Cellular deconvolution analysis revealed differences in immune and stromal cell proportions between GAC and PTT, as shown in Figure 5A. Among the identified cell types, B cells, macrophages, neutrophils, mast cells, and cancer-associated fibroblasts (CAFs) exhibited notable shifts in abundance across module groups.

**Figure 5.**
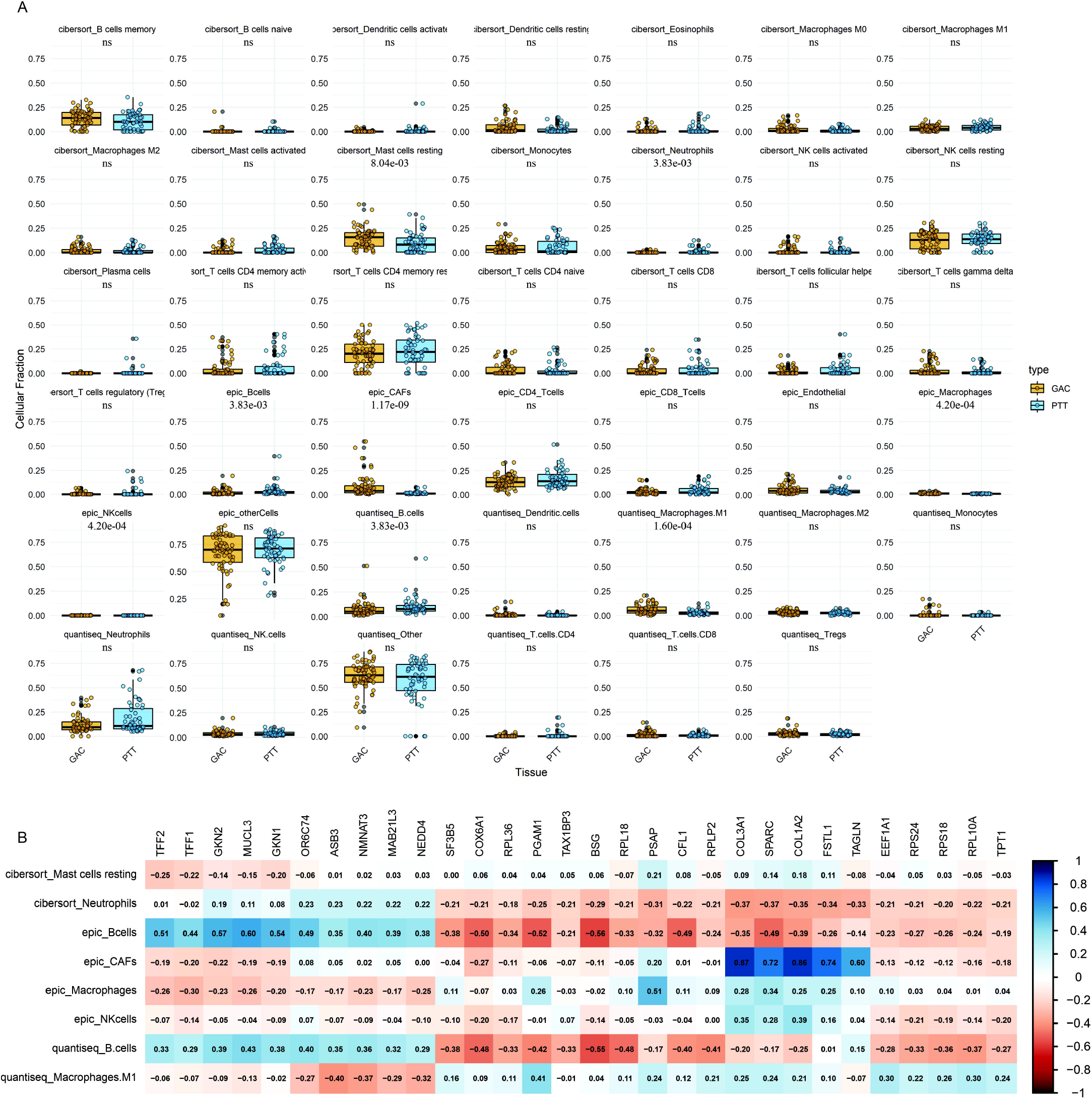
Cellular Deconvolution and Hub Genes. (A) Average differences in immune cell composition between GAC and PTT groups; (B) Associations between immune cell proportions and the expression of immune-related genes in GAC samples.

Correlation heatmaps (Figure 5B) indicate specific relationships between immune cell populations and the expression of hub genes. Genes from the MEblack module, particularly *TFF2, TFF1, GKN2, MUCL3*, and *GKN1*, displayed consistent positive correlations with B cells (correlations between 0.29 and 0.61) across multiple estimation methods (e.g., EPIC, quantiseq). These same genes showed negative correlations with neutrophils and mast cells, with values ranging from −0.21 to −0.45. MEturquoise hub genes, such as *COL1A2, COL3A1, TAGLN*, and *SPARC*, were positively associated with CAFs (up to r = 0.72), suggesting alignment with the fibroblast-rich tumor stroma. Genes like *EEF1A1, RPS18*, and *PGAM1* showed negative correlations with B cells, and more variable correlations with macrophages and NK cells, generally not exceeding |0.30|.

Different modules showed distinct correlation patterns with immune and stromal components. MEblack genes tend to co-vary with adaptive immune elements, particularly B cells, while MEturquoise genes are more closely associated with fibroblastic or innate immune components. MEblue genes presented more heterogeneous correlations, generally weaker in magnitude and less consistent across cell types.

## Discussion

Facing the complexities of GAC, there is a need to develop advanced molecular biomarkers that can refine both diagnosis and treatment, ultimately enhancing clinical outcomes. Our analysis of transcriptomic comparison between GAC tissues and their paired peritumoral PTT highlights two key biological aspects: the heterogeneity between patients and a continuum of molecular transformation. In the unsupervised clustering analysis (Figure 1A), the GAC and PTT samples do not clearly separate but instead form mixed clusters. This tissue overlap suggests that inter-individual variability may be greater than the average differences between the tumor and peritumoral tissues. Hub genes in the MEbrown, MEgreen and MEturquoise modules have higher expressions in two of the three clusters created by the unsupervised analysis in heatmap (from left to right, cluster 1 and 2); while MEmagenta, MEblack and MEblue are more characteristic of individual clusters, cluster 1, 2 and 3 respectively. The pattern identified also supports the concept of the cancerization field, in which histologically “normal PTTs” already exhibits early molecular changes associated with neoplastic transformation.

When the impact of inter-individual differences was mitigated by implementing paired analysis (see Figure 2B), more consistent disparities in the expression of core genes became evident. This difference is evidenced in the hub genes present in MEblack and MEmagenta (fifth and first lines of figure 4B, respectively), which represent the two groups of genes with the most significant disparities in expression between GAC and PTT. Thus, due to the greater expression and correlation of the MEblack genes with PTT, and of MEmagenta with GAC, we hypothesized that MEblack may be a signature of a normal mucosa and MEmagenta of a fully transformed cancerous epithelium.

Additionally, we investigated the associations between clinical features and the hub genes that exhibited the most pronounced differences in expression between GAC and PTT. Notably, *TFF1, TFF2, GKN1, GKN2*, and *MUCL3* - hub genes of the MEblack module - exhibited consistent positive correlations with MMR status, hPSM2, and hMLH1 expression, reinforcing their potential involvement in the maintenance of normal mucosa by genomic stability. Their inverse correlations with staging variables, although modest, continue to suggest their relevance to early-stage or less aggressive tumor contexts. On the other hand, genes from the MEmagenta module stand out for their positive correlations with clinical progression metrics, including ypTNM, T, and ypN. This reinforces their potential role as markers of tumor burden or advanced disease. MEblue genes such as *ASB3, NMNAT3*, and *MAB21L3* display a more heterogeneous correlation profile. They show moderate positive associations with PDL1 and weaker or inconsistent correlations with staging or differentiation variables. Unlike MEblack, MEmagenta and MEblue hub genes appear tightly connected to tumor expansion dynamics. Thus, the current findings emphasize that MEblack, MEmagenta and MEblue, represent transcriptional programs with distinct clinical implications: epithelial integrity, tumor burden and plasticity respectively.

Exploring functional enrichment of the three more biologically relevant modules cited above, we observed that the genes present in MEmagenta are related to the organization of the extracellular matrix (ECM). The ECM provides structural support and influences cell behavior, such as the proliferation and migration of cells in cancer (Pickup et al. 2014). In this context, the five hub genes identified in this module, such as *COL3A1*, which encodes type III collagen, and *SPARC*, which modulates cell-matrix interactions and tissue remodeling, were overexpressed in our GAC samples, as corroborated by the literature (Zhou et al. 2022). *COL1A2* is involved in type I collagen production and its high expression, as in out GAC tissues, is related to gastric tumor size and depth of invasion (Li et al. 2016), *FSTL1*, which regulates cell growth and angiogenesis(Wu et al. 2021), also plays a key role in tumor progression, while *TAGLN* is associated with cell motility and invasiveness (Zhou et al. 2016).

GC progression involves disruptions in epithelial maintenance, digestion, and xenobiotic responses. Loss of epithelial integrity, metabolic imbalance, and dysregulated xenobiotic response pathways, especially those involved in microbial sensing, can amplify local inflammation and immune evasion (Faubert et al. 2020). In the MEblack module, hub genes including *GKN1, GKN2 TFF1* and *TFF2*(Chung Nien Chin et al. 2020) are intimately involved in the preservation of gastric mucosal homeostasis. These genes contribute to prevent damage induced by gastric acid and digestive enzymes, thus preserving the functional integrity of the digestive environment. Importantly, all five genes were found to be downregulated in our GAC samples, suggesting a coordinated loss of mucosal defense and regenerative capacity during tumor progression. *GKN2* and *TFF1* may exert synergistic antiproliferative and pro-apoptotic effects, while *MUCL3* supports epithelial-microbial interface integrity, linking digestion, inflammation, and tumor biology (Arai et al. 2024).

The MEblue module exhibited significant enrichment in GO terms related to cell-cell interactions and structural integrity, indicating its involvement in critical cellular processes. Among its hub genes, *NMNAT3* is known for its role in NAD□ biosynthesis—a pathway essential for cellular metabolism and stress adaptation—and has been associated with cancer cell survival through p53 modulation (Liu et al. 2021). *OR6C74*, an olfactory receptor gene, may influence cell signaling, although its role in oncogenesis remains poorly defined (Chung et al. 2022). *MAB21L3* lacks consistent evidence of association with cancer, and its biological functions are still under investigation (Qin et al. 2024). Finally, *NEDD4* is a well-known E3 ubiquitin-protein ligase, frequently implicated in cancer due to its role in the degradation of tumor suppressor proteins (Sun et al. 2014). However, despite their known or putative roles in other malignancies, the specific contribution of these genes to GC remains unclear, underscoring the need for further functional studies to elucidate their roles within this tumor context.

The genes in the *TFF1, TFF2, GKN2, MUCL3*, and *GKN1* (MEblack) show a consistently positive correlation with the fraction of B cells GAC and a moderately negative correlation with resting mast cells and neutrophils. This pattern suggests that, among the findings from the cellular deconvolution analysis, there is an association between the expression of these epithelial factors and the presence of B lymphocytes, whereas a higher abundance of mast cells and neutrophils is linked to reduced expression levels of these genes. It is important to emphasize that these relationships are co□variation observations and do not permit any claims of causality or direct regulatory mechanisms between the genes and each cell type.

On the other hand, the (*COL1A2, COL3A1, TAGLN*, and *SPARC* (MEturquoise) is strongly associated with CAFs and shows a negative correlation with the fraction of B cells. This finding may indicates that tumor regions with greater CAF activity - reflected by high levels of collagen and extracellular matrix proteins - tend to have lower B□cell infiltration, possibly due to the physical□chemical barriers created by the extracellular matrix or to pro□fibroblastic signaling. Even so, the observed correlations do not prove that CAFs directly prevent B□cell recruitment or explain all variations in gene expression, but they do reveal distinct cellular□composition profiles that can guide future investigations.

Overall, our findings show that the identified coexpression modules correspond to distinct biological features of gastric adenocarcinoma versus peritumoral tissue. The MEblack module (*TFF1, TFF2, GKN1, GKN2, MUCL3*) is associated with a more intact mucosal state and higher B-cell presence in GAC compared to PTT, suggesting these factors help maintain the gastric epithelium and genomic integrity. In contrast, the MEturquoise module (*COL1A2, COL3A1, TAGLN, SPARC*) comprises genes linked to CAF enrichment and collagen deposition, which inversely correlate with B-cell infiltration, without definitively establishing stromal stiffness as the cause of lymphocyte exclusion. The MEmagenta module highlights extracellular matrix genes tied to tumor size and invasion, while MEblue contains genes with more variable, as-yet-uncharacterized correlations in GAC. Together, these profiles suggest a possible molecular continuum between PTT and GAC.

However, certain limitations may impact the interpretation of the data. First, the analysis was conducted with a considerable but limited number of samples. Additionally, although we identified hub genes with high relevance, functional validation of these genes is still necessary. Therefore, while our findings provide a good foundation for further single cell analysis, future investigations must address these limitations and explore the functional implications of the identified genes in more depth.

## Acknowledgments

The authors thank CAPES (Coordenação de Aperfeiçoamento de Pessoal de Nível Superior) for providing a doctoral scholarship to R. M. da S. Mourão. We also thank CCAD (Centro de Computação de Alto Desempenho da Universidade Federal do Pará - High Performance Computing Center) for their support in this research. Additionally, we acknowledge the Fundação Amazônia de Amparo a Estudos e Pesquisas (Fapespa) for funding this study.

## Competing interests

The authors declare no competing interests.

## Author contributions

R.M.S.M. and F.C.M. conceived the statistical analysis of the data, interpretation of the results, and drafting of the manuscript. J.M.C.S., D.S.A.C., and V.C.S.S. assisted in data processing and revision of the manuscript. A.F.V., L.L.M. provided technical assistance in sample processing and sequencing. A.K.M.A., S.D., and W.F. B. obtained and reviewed the patients’ clinical and pathological data. P.P.A. and F.C.M. contributed to the conceptualization of the article and funding the acquisition. R.M.S.M. and F.C.M. are the first authors of this work. All authors reviewed and approved the final version of the manuscript.

## Supplementary Material: https://shre.ink/supplementary-materials (Link to Tables only)

**Table S1. Normalized gene expression. Gene expression values for each sample based on transcript quantification. Rows correspond to ind**ividual samples, and columns represent genes. The matrix includes expression levels used in downstream co-expression network analyses.

**Table S2. GS and MM**. Correlation metrics between gene expression and clinical traits. For each gene, Gene Significance (GS) indicates the strength of association with a given trait, while Module Membership (MM) measures the correlation of the gene with its respective co-expression module. These values help identify hub genes with potential biological relevance.

**Table S3. Ontology**. Results of Gene Ontology enrichment analysis for selected gene sets. Each row corresponds to an enriched GO term and includes statistical parameters such as adjusted p-values (p.adjust), enrichment scores and associated gene lists. These annotations help infer biological processes overrepresented in the dataset.

**Table S4. AUC**. Performance evaluation of gene-based classifiers or predictive models using the AUC. Higher AUC values indicate better discriminatory power, useful for identifying genes or signatures with biomarker potential.

**Figure S1. Principal Component Analysis (PCA) differentiating GAC and PTT samples**. Each point represents an individual sample, plotted based on the first two principal components (PC1 and PC2). Samples are color-coded according to their respective tumor groups. The distinct clustering of GAC and PTT samples indicates substantial differences in their molecular characteristics.

## Notes

### Competing Interest Statement

The authors have declared no competing interest.

