## Supplementary figures and images for "EXPRESSION INSIGHTS INTO GASTRIC ADENOCARCINOMA: NETWORK ANALYSIS REVEALS KEY HUB GENES AND FUNCTIONAL MODULES"

### S1. PCA.tiff

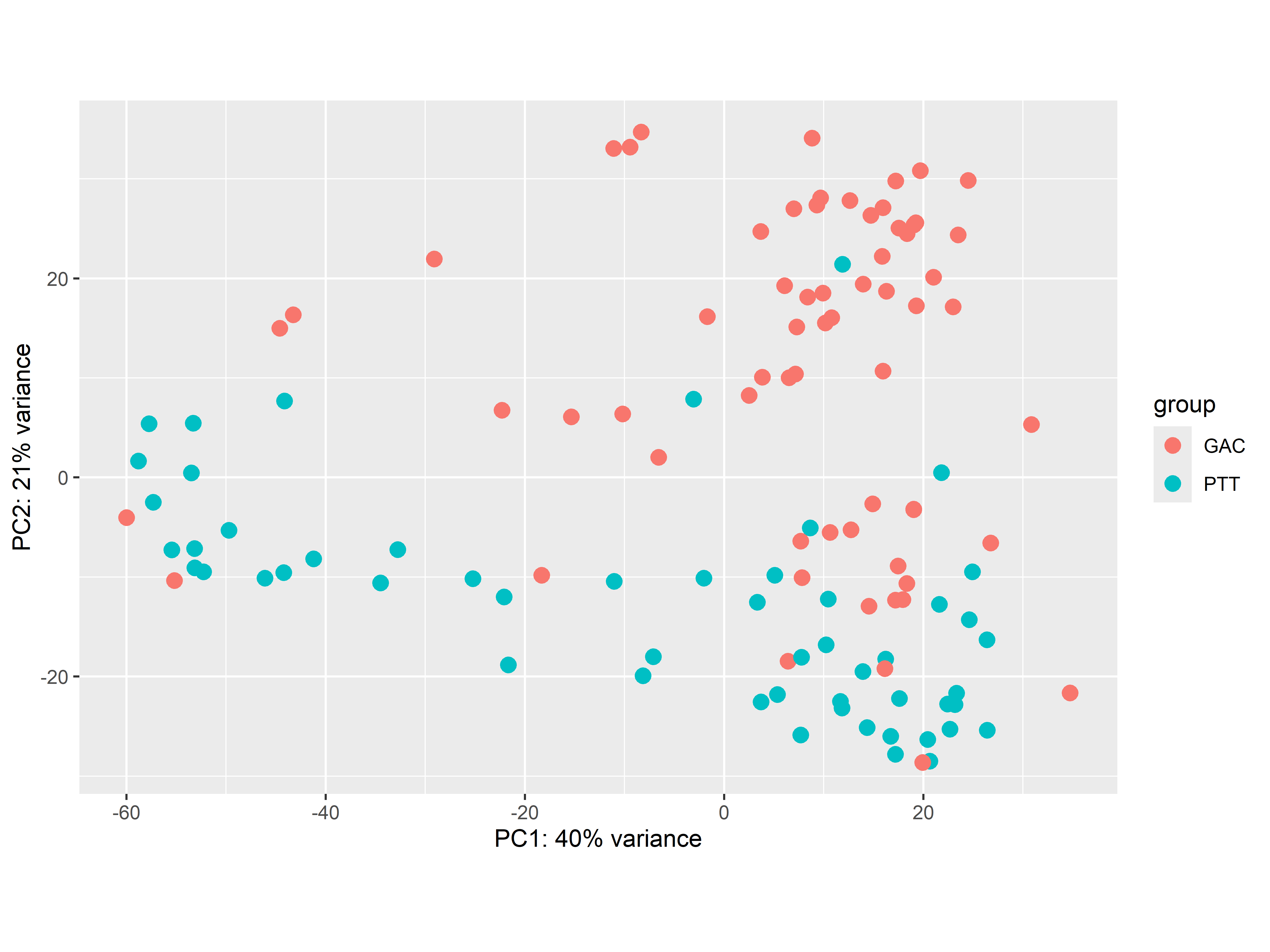
